# RNA viruses in water shape the viromes of shrimp and fish in aquaculture systems

**DOI:** 10.1101/2025.10.19.683331

**Authors:** Dandan Zhou, Chengguang Xing, Shanshan Liu, Zhi Liu, Yixuan Liu, Yinqing Wu, Hualong Su, Danrong Xian, Guangyu Guo, Jiarong Cheng, Menghuang Xu, Man Chen, Peiyun Han, Chengcheng Wu, Jiamin Zeng, Xinyi He, Jianhu Pang, Qiang Lu, Chao Zhang, Taiyu Wang, Cancan Zhang, Yichun Xu, Xiaocheng Tang, Shuqiao Zhang, Zhixuan Deng, Shenzheng Zeng, Shaoping Weng, Mang Shi, Jianguo He

## Abstract

Aquatic environments host immense RNA viral diversity, yet the extent to which waterborne viruses influence the virome of aquatic animals remains unclear. Here, we conducted a comprehensive longitudinal metatranscriptomic analysis of water, shrimp, and fish across multiple culture stages in aquaculture systems. We identified 3,211 RNA viruses spanning 21 viral supergroups, with 97.4% representing novel species. Water exhibited the greatest viral diversity (3,093 species), followed by shrimp (1,280) and fish (398), with most viruses detected in shrimp and fish also detected in water. Diversity analyses revealed increasing richness and significant compositional shifts in 92 high-abundance viruses (>1,000 RPM) across culture stages. Viral abundance in aquatic animals was strongly correlated with that in water, and the temporal trajectories of the 16 most abundant viruses were closely synchronized between water and shrimp. Furthermore, 337 virus species were shared among shrimp and two fish species, of which 309 were present in water. These findings demonstrated that RNA viruses in water shape the composition and dynamics of aquatic animal viromes, underscoring the importance of water virome surveillance and management for disease prevention in shrimp aquaculture.

## Introduction

Aquatic environments serve as important reservoirs of viruses^1–3^, and large-scale discovery efforts have unveiled remarkable diversity of RNA viruses across diverse aquatic habitats^4–9^. For example, metagenomic analyses of water samples from Yangshan Deep-Water Harbor near Shanghai has doubled the known diversity of RNA viruses^10^, while abundant novel RNA viruses have also been reported in alpine lake on the Tibetan Plateau^11^. Collectively, these studies highlight water as a primary source of RNA viruses and underscore its central role in shaping global viral diversity. Beyond their diversity, aquatic viruses are now recognized as major regulators of ecological and evolutionary processes in aquatic ecosystems^12–14^. Previous study suggested potential links between virus and marine carbon cycling, proposing that viral infections of aquatic hosts may influence the biological pump^15^. Moreover, aquatic viruses are thought to encode auxiliary metabolic genes involved in diverse host metabolisms, and might predict marine carbon export^16^, pointing to their role in host metabolism and ecosystem functioning.

Importantly, virome studies of aquatic animals have revealed extensive viral diversity, including viruses affecting economically important animals^17–21^. In addition, some studies have suggested a potential connection between host virome and their environments. For instance, oysters have been shown to share RNA viruses with other mollusks and surrounding seawater, suggesting strong connectivity of viral communities between aquatic species and their environments^22^. Another study has showed differences in viral composition between cultured and wild European brown shrimp, indicating that certain viruses may become more abundant under sub-optimal aquaculture conditions^23^. However, despite progress in characterizing aquatic RNA virus diversity, the direct influence of waterborne viruses on the virome composition of aquatic animals remains poorly understood.

This knowledge gap is particularly important for aquaculture. The Pacific white shrimp (*Litopenaeus vannamei*), the most widely farmed crustacean globally, is severely threatened by viral disease outbreaks that cause substantial economic losses^24^. Water quality is a critical determinant of shrimp health^25,26^: aquaculture systems using groundwater or disinfected recirculating water exhibit lower pathogen loads and fewer outbreaks compared with systems relying on untreated water^27^. Similarly, coastal areas with better water quality are generally more favorable for shrimp farming, while higher organic loading is associated with increased disease risks^28^.

Building upon these previous findings, the present study conducted a comprehensive and well-designed virome analysis of water, shrimp, and fish across different culture stages in shrimp aquaculture systems. We characterized total viromes to assess the composition, diversity, and temporal dynamics of high-abundance viruses, examined the contribution of waterborne viruses to the virome of shrimp and fish, and explored virus sharing across hosts. Our results demonstrated that RNA viruses present in water substantially shape the virome composition of aquaculture animals, providing new insights into disease ecology and informing strategies for prevention and control in shrimp aquaculture.

## Results

### Study design and sampling strategy

From June to September 2020, we conducted a longitudinal sampling of shrimp aquaculture ponds operating under two distinct culture systems: shrimp monoculture and shrimp-fish polyculture systems in Maoming, Guangdong Province, China (Fig. S1). Sampling covered an entire shrimp culture cycle and included five key time points: prior to larval stocking (Stage 0) and four subsequent stages representing early (Stage 1 and Stage 2), middle (Stage 3), and late (Stage 4) phases of cultivation. In total, 525 Pacific white shrimp (*Litopenaeus vannamei*), 60 tilapia (*Oreochromis niloticus × O. aureus*), 60 red drum (*Sciaenops ocellatus*), and 1,050 liters of water were collected. These samples were categorized by sample type (shrimp, tilapia, red drum, and water), culture system (monoculture or polyculture), and culture stage (Stage 0-4), resulting in 31 pooled shrimp samples, 6 pooled tilapia samples, 6 pooled red drum samples, and 41 pooled water samples for subsequent RNA sequencing and virome profiling (Table S1).

### Characterization of the RNA viromes

RNA sequencing of 84 pooled samples yielded an average of 66 million clean reads per library, which were assembled into 2.66 million contigs (> 600 bp) (Table S1). After merging and clustering, 1.66 million non-redundant contigs were screened for viral sequences, yielding 3,211 RNA virus contigs ranging from 746 to 18,112 bp in length, all identified based on the presence of RNA-dependent RNA polymerase (RdRP) gene (Table S2). Phylogenetic analyses assigned 3,205 viruses to 21 known “supergroups” (corresponding to recognized viral orders or classes)^29^, while six remained unclassified (Fig. S2 and Table S2). Based on RdRP protein and genome identity thresholds, 97.4% (n = 3,128) represented putative novel viral species, highlighting substantial undiscovered diversity within shrimp aquaculture systems.

Viral abundance and richness varied markedly across sample type (Table S3 and S4). Shrimp samples exhibited the highest viral abundance (median: 633,604 RPM, range: 17,715 to 927,312 RPM), followed by water (median: 453,681 RPM, range: 14,276 to 915,291 RPM) and fish (median: 8,293 RPM, range: 1.7 to 71,215 RPM) (Table S1). In terms of richness, water samples harbored the greatest number of viral species (n = 3,093), followed by shrimp (1,280), and fish (398 in total, 300 in tilapia and 304 in red drum) (Fig. 1a). Across all hosts, the most diverse viral supergroups were Picorna-Calici (number of viral species n = 925), Tombus-Noda (n = 637), and Narna-Levi (n = 487), with consistent patterns among water, shrimp, and fish (Fig. 1a and Table S4). In terms of relative abundance, the Picorna-Calici supergroup dominated all viromes (median 95.31%, range 9.78% to 100% of viral RNA reads), followed by Hepe-Virga (median 0.15%, range 0 to 72.37%) and Narna-Levi (median 0.36%, range 0 to 53.87%) (Fig. 1b and Table S5).

**Fig. 1.**
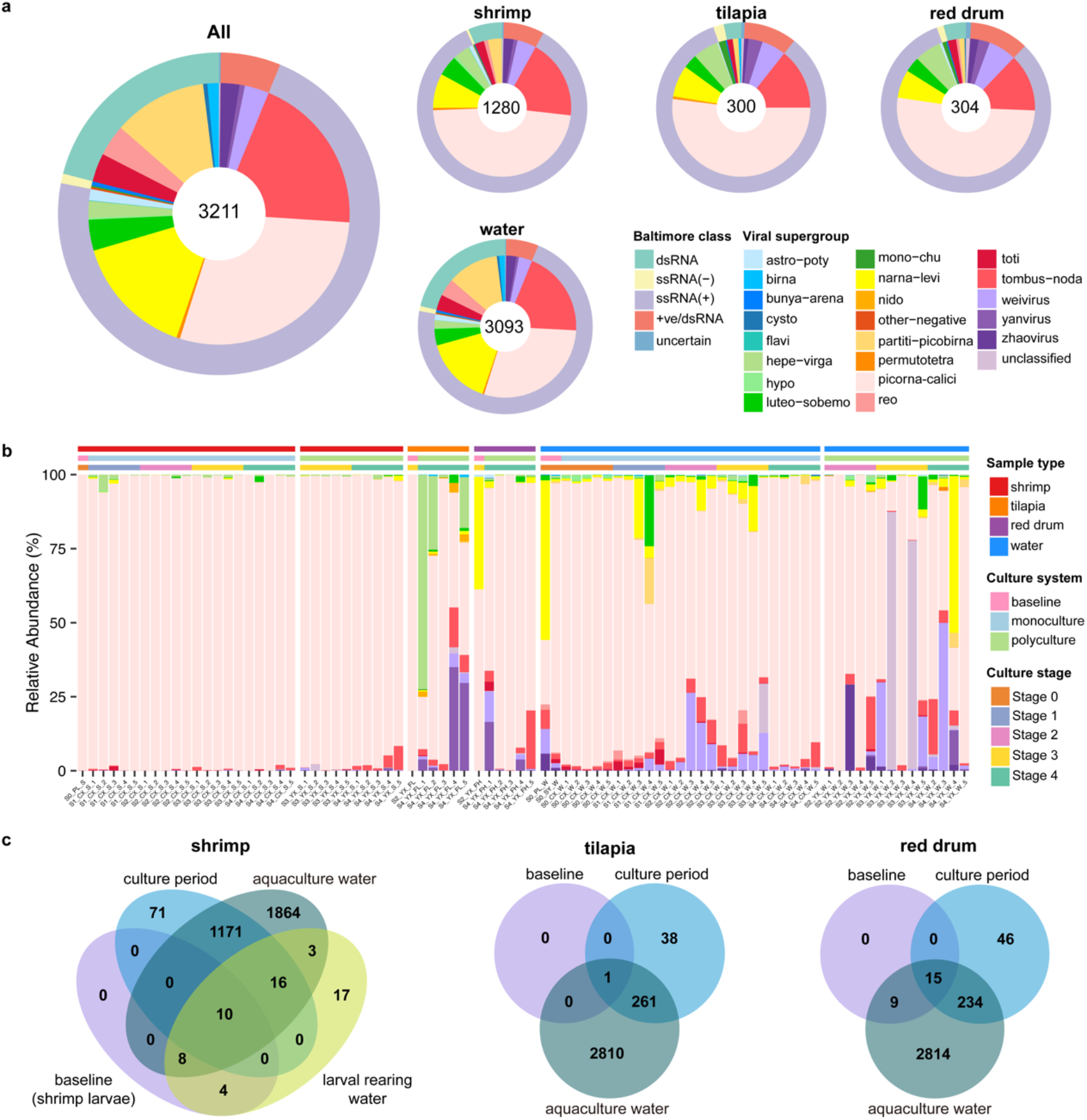
Overview of the RNA viruses identified in shrimp aquaculture systems. **a**, Number of virus species categorized by supergroups and baltimore classes across shrimp, tilapia, red drum, and water. **b**, Relative abundance of viral supergroups in each metatranscriptomics library. **c**, Venn diagrams showing the overlap of virus species between water and each host organism (shrimp, tilapia, and red drum).

Temporally, only a few viruses were detected in baseline samples (22 in shrimp larvae, 1–24 in fish), whereas the majority emerged during the culture period (1268 in shrimp, 295–300 in fish). Notably, 77–92.7% of these viruses overlapped with those found in water, suggesting that water serves as a major source and reservoir of RNA viruses in aquaculture systems (Fig. 1c).

### Diversity and composition of high-abundance viruses

To identify likely shrimp-associated viruses, we focused on those with high abundance (≥ 1000 RPM) in shrimp samples^30^. A total of 92 such viral species meeting this criterion were identified and subsequently validated by RT-PCR and Sanger sequencing. These viruses belonged to eight major RNA virus supergroups—Picorna-Calici (65), Hepe-Virga (8), Tombus-Noda (8), Luteo-Sobemo (3), Narna-Levi (3), Weivirus (2), Partiti-Picobirna (1), and Yanvirus (1) —plus one unclassified virus (Table S2). Collectively, these viruses accounted for 87.2%–99.9% of total viral RNA reads in shrimp and also contributed notably to the viromes of tilapia (11.2%–100%), red drum (13.5%–99.8%), and water (0.06%–95.6%) (Table S6). Their widespread presence and high prevalence across shrimp and water suggest they are common and ecologically relevant members of the virome in shrimp aquaculture systems (Fig. S3).

To assess viral diversity, we calculated total viral abundance, species richness, and Shannon index (Table S7). These metrics varied significantly among sample types (Kruskal-Wallis test; *P* < 0.05 for richness, *P* < 0.001 for abundance and Shannon index; Fig. 2a). Shrimp showed the highest viral abundance but significant lower Shannon index, potentially reflecting infection by dominant viruses. The viral species richness in shrimp increased progressively across culture stages (Kruskal-Wallis test, *P* < 0.01), although abundance and Shannon index remained stable (Fig. 2b). Fish exhibited marked increases in both species richness and abundance in Stage 4 compared to baseline (Fig. 2c). Water also showed significant stage-dependent variation in species richness (Kruskal-Wallis test, *P* < 0.001), but no consistent trends in shrimp or water were associated with culture systems (Fig. 2b & d).

**Fig. 2.**
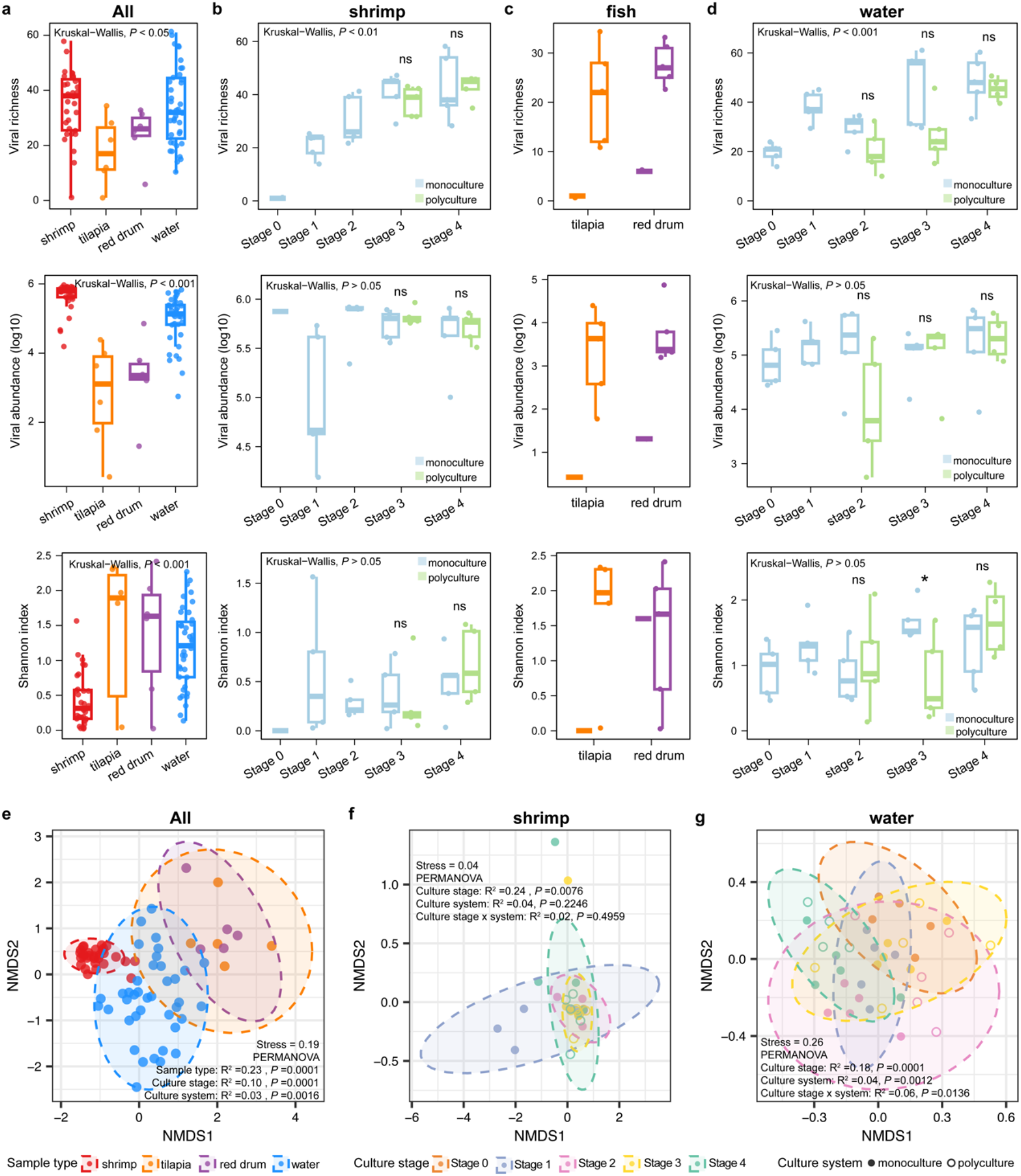
Diversity patterns of high-abundance viruses across sample types, culture systems, and culture stages. **a**, Alpha diversity indexes (viral species richness, viral abundance, and Shannon index) across sample types. **b**, Alpha diversity in shrimp across culture systems and culture stages. Asterisks indicate statistical significance (t-test, ns, *P* > 0.05; *, *P* < 0.05). **c**, Alpha diversity in tilapia and red drum across culture stages. **d**, Alpha diversity in water across culture systems and culture stages. **e**, NMDS plot of viral composition among sample types. *P*-values from PERMANOVA are shown. **f**, NMDS plot of viral composition in shrimp across culture stages. **g**, NMDS plot of viral composition in water across culture stages.

Non-metric multidimensional scaling (NMDS) revealed distinct viral compositions among shrimp, fish and water (Fig. 2e). PERMANOVA analyses confirmed that sample type was the primary driver of variation (R^2^ = 0.23, *P* = 0.0001), followed by culture stage (R^2^ = 0.10, P = 0.0001) and culture system (R^2^ = 0.03, *P* = 0.0016) (Fig. 2e). Viral composition in shrimp was significantly shaped by culture stage (R^2^ = 0.24, *P* = 0.0076) (Fig. 2f), while viral composition in water was significantly influenced by both in culture stage and culture system (R^2^ = 0.18 and R^2^ = 0.04, respectively, Fig. 2g).

### Temporal dynamics of dominant virus species

We next examined the temporal dynamics of the 16 most abundant viruses (each >10,000 RPM), which collectively accounted for the majority of viral RNA reads in shrimp and water. Overall, viral abundance increased progressively across culture stages in both shrimp and water, with fish showing much lower levels, indicating strong host specificity (Fig. 3a). Despite variation among individual species, four general temporal patterns were observed: (i) persistent high levels (n = 1, e.g. Dianbai picorna-calici-like virus 24), (ii) gradual increases (n=1, e.g. Dianbai picorna-calici-like virus 686), (iii) mid-stage peaks (n=4, e.g. Dianbai picorna-calici-like virus 191 and Dianbai picorna-calici-like virus 377), or (iv) late-stage surges (n=7, e.g. Dianbai picorna-calici-like virus 115 and Dianbai tombus-noda-like virus 2) (Fig. 3b).

**Fig. 3.**
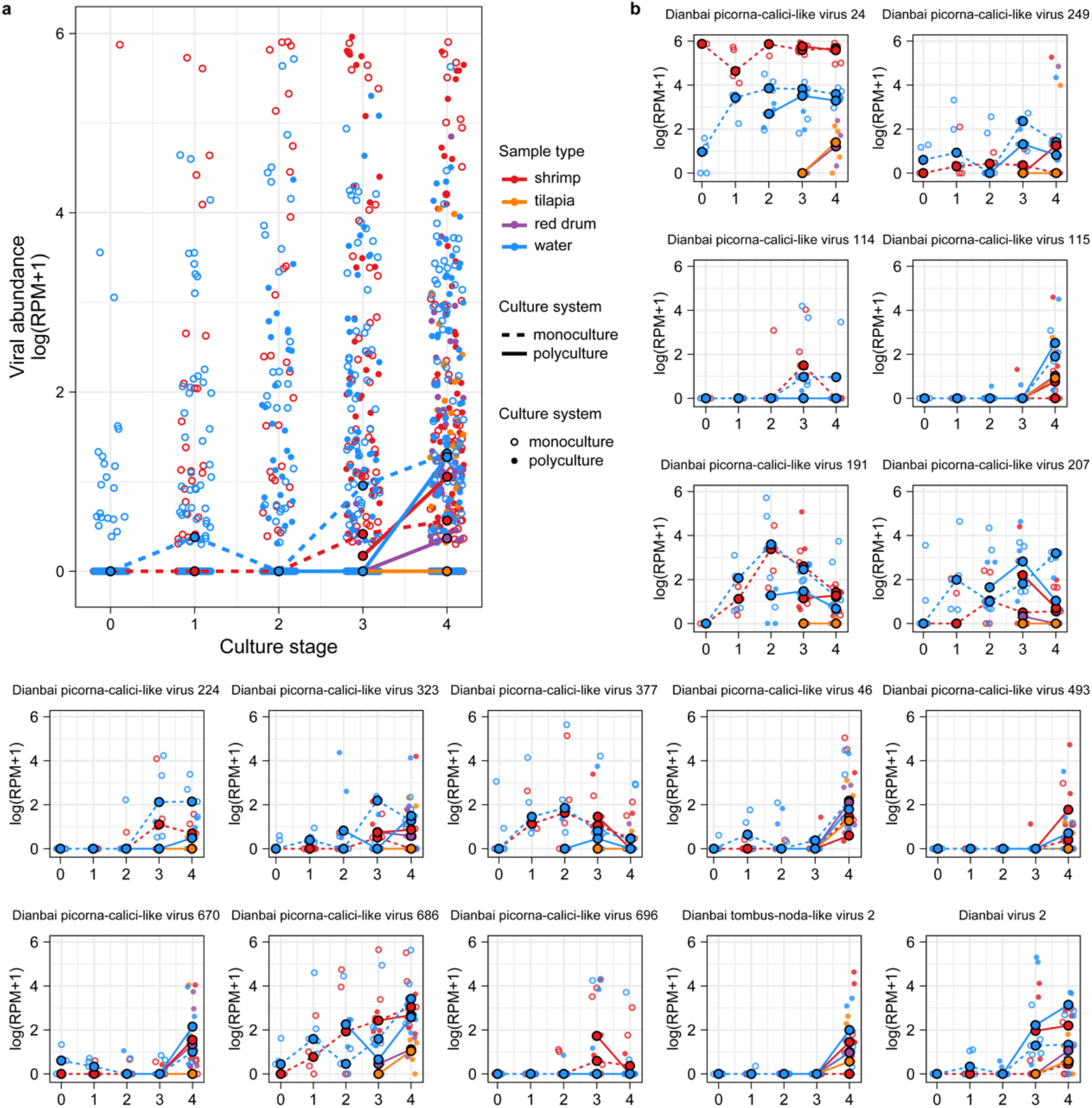
Temporal dynamics of dominant virus species. **a**, Temporal changes in the top 16 high-abundance viruses across culture stages. The x-axis show culture stages, and the y-axis denotes log-transformed viral abundance (RPM). Hollow and solid circles represent viral abundance in monoculture samples, respectively, with each point corresponding to a sample. Median values across samples for each virus are connected by lines—dashed for monoculture, solid for polyculture. Sample types are color-coded: red (shrimp), orange (tilapia), purple (red drum), and blue (water). **b**, Temporal abundance profiles of individual viral species, ranked by their maximum abundance in shrimp. Each plot shows the median abundance across culture stages for a single virus.

Notably, shrimp and water showed similar dynamic trends, suggesting that aquatic viral dynamics are closely coupled with changes in the shrimp virome. Viral abundance patterns also differed between culture systems: several picorna-calici-like viruses (e.g., Dianbai picorna-calici-like virus 24, Dianbai picorna-calici-like virus 249, and Dianbai picorna-calici-like virus 377) were enriched in the monoculture system, while Dianbai tombus-noda-like virus 2 and Dianbai virus 2 predominated in polyculture systems (Fig. 3b).

### Correlation of viral abundance between water and aquatic animals

To assess the relationship between viral abundance in water and those in aquatic animals, we performed correlation analyses across sample types. Viral abundance in water was strongly and positively correlated with that in shrimp under both culture system (Spearman’s R = 0.59 and 0.64 for monoculture and polyculture, respectively; *P* < 2.2e^-16^) (Fig. 4a). Notably, the stronger correlation coefficient observed in polyculture system suggests enhanced viral exchange or shared exposure among co-cultured species. Similarly, significant positive correlations were detected for tilapia (R = 0.51) and red drum (R = 0.63) (Fig. 4b & c). A complementary analysis incorporating all 3,211 detected viral species yielded consistent results (Fig. S4), reinforcing the notion that water acts as a central reservoir shaping the virome of aquatic animals.

**Fig. 4.**
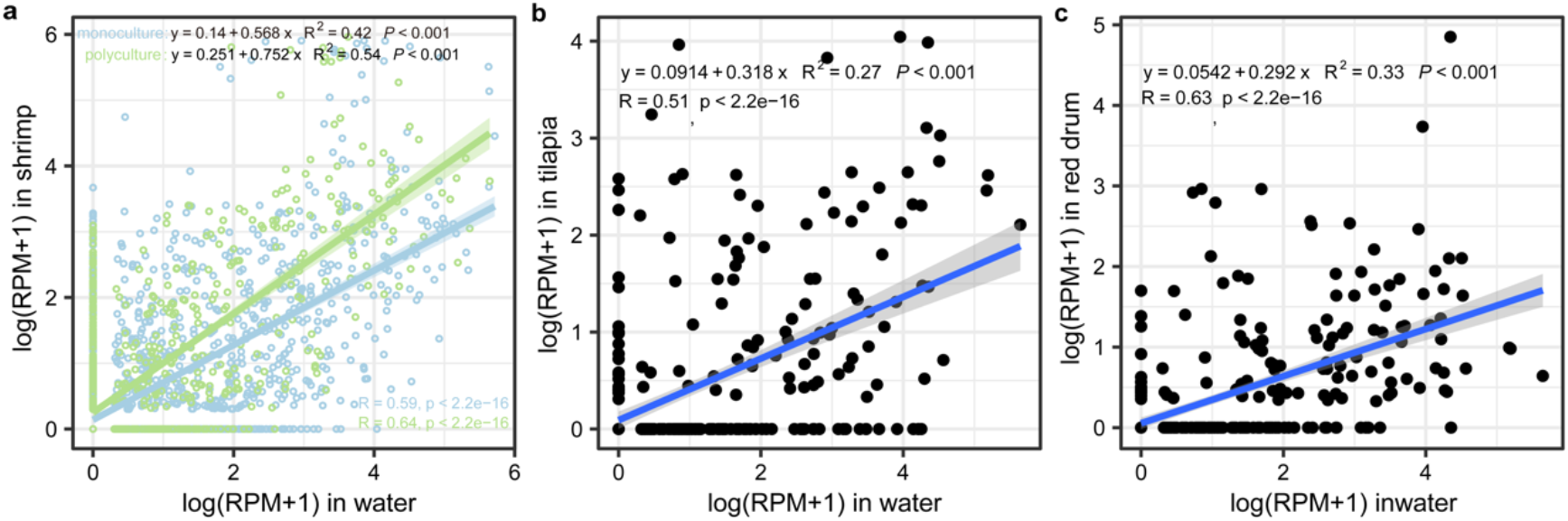
Correlation of high-abundance virus loads between aquatic animals and water. **a**, Correlation of viral abundance between shrimp and water. Shaded areas represent 95% confidence intervals. **b**, Correlation between tilapia and water. **c**, Correlation between red drum and water.

### Viral sharing among shrimp and fish

We next examined viral sharing among shrimp, tilapia, and red drum. A total of 337 virus species (10.5%) were shared among the three host species, with 179 detected in all three (Fig. 5a & b and Fig. S5). Before farming, no viruses were shared between shrimp and fish, whereas during the culture period, 332 viral species became shared (Fig. S6a & b). Remarkably, 91.7% (309/337) of these shared viruses were also detected in water (Fig. 5a & b), underscoring its role as a key conduit for viral exchange between species. The number of shared viruses in shrimp increased progressively over culture stages, and shrimp in polyculture system harbored more shared viruses with fish (283 species, 84%) than those in the monoculture system (251 species, 74.5%), especially during Stage 3–4 (Fig. 5c and Fig. S6b-d). Shared viruses also had significantly higher prevalence in shrimp from polyculture system (*P* < 0.001), indicating that co-cultivation facilitates cross-species viral transmission (Fig. 5d).

**Fig. 5.**
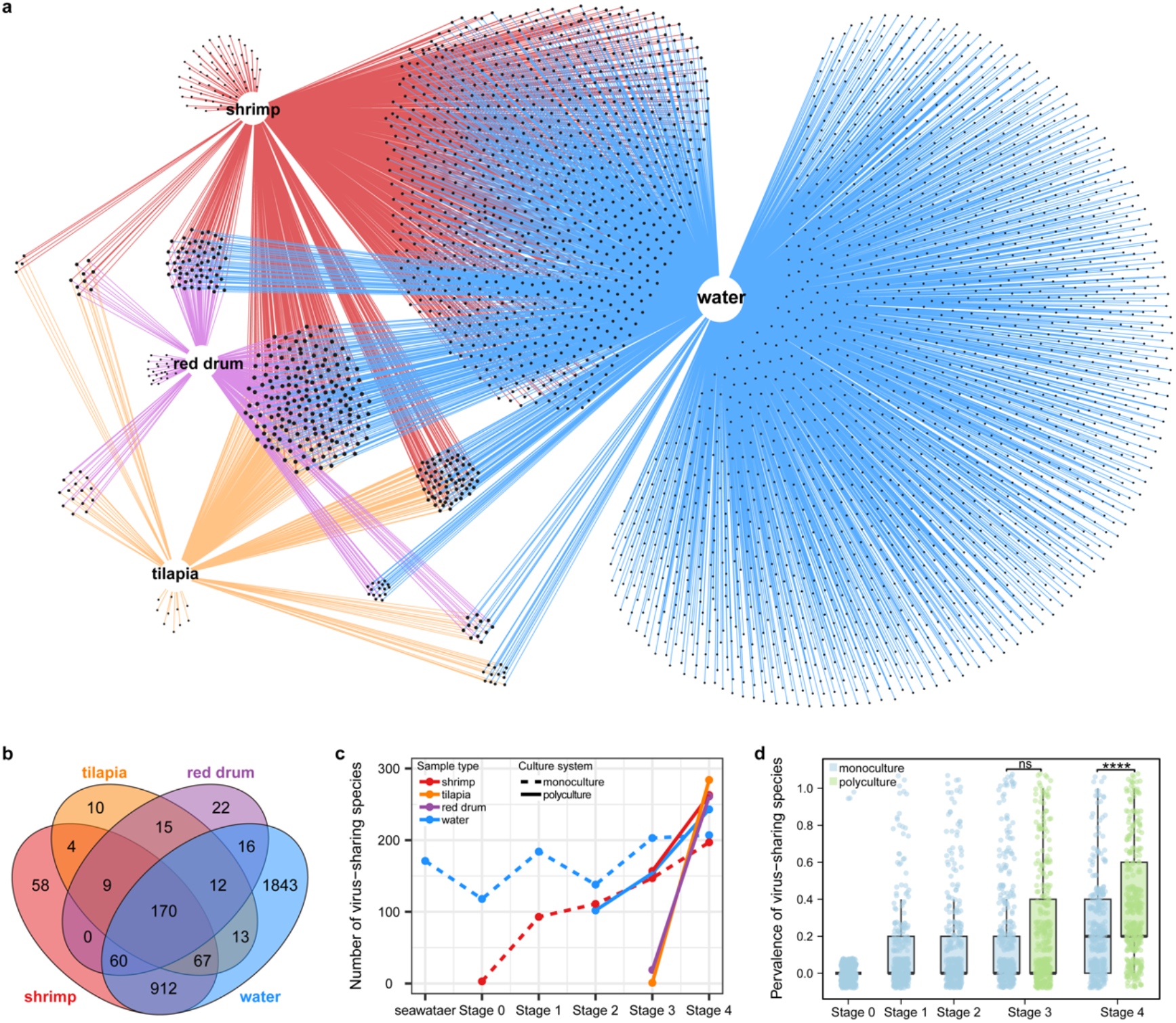
Virus sharing among shrimp, fish, and water. **a**, Network visualization of viral sharing among shrimp, fish and water. **b**, Venn diagram showing shared and unique virus species among shrimp, tilapia, red drum, and water. **c**, Number of shared virus species across culture stages for each sample type. **d**, Comparison of the prevalence of shared viruses across culture stages and culture systems. Asterisks indicate statistical significance (t-test; ns, P > 0.05; ****, P < 0.0001).

## Discussion

This study provides a comprehensive characterization of RNA viromes in shrimp monoculture and shrimp–fish polyculture systems. We uncovered a remarkable RNA viral diversity in both aquatic animals and their surrounding water, revealing that water functions as a major viral reservoir. Indeed, most viruses detected in shrimp and fish originated from water, and numerous species were shared among these hosts, emphasizing the central role of water viromes in shaping aquatic animal virome composition and driving cross-species viral transmission.

Aquatic environments are well recognized as vast viral reservoirs that sustain extraordinary RNA virus diversity across taxa^1^. The high viral diversity in water likely reflects its ecological complexity, harboring diverse aquatic animals, plants, and microbes that together facilitate viral persistence and exchange^31^. In our aquaculture systems, we identified 3,093 RNA viruses in water,1280 in shrimp and 398 in fish— spanning a wide range of taxonomic groups and largely comprising novel species. This scale of discovery aligns with previous observations from marine and freshwater environments and reinforces the view that aquatic viromes are a central component of viral evolution and ecology5,11. Importantly, the virome composition in shrimp and fish expanded dramatically after introduction into aquaculture systems, with the majority of these viruses (92.7% in shrimp, 87% in tilapia, and 77% in red drum) also present in water. This strongly supports the notion that water virome serve as the primary source shaping viral diversity in aquatic animals.

Cross-species viral transmission is a key evolutionary process enabling viruses to overcome host barriers and occasionally trigger disease emergence^32^. In our study, 337 virus species were shared among shrimp, tilapia, and red drum—none of which were shared prior to farming. This striking shift indicates extensive cross-species transmission during aquaculture. The fact that 91.7% of these shared viruses were also detected in water suggests that water acts as an efficient medium for viral exchange. Unlike terrestrial systems, where blood-feeding arthropods or plant-feeding insects mediate transmission^33^, aquatic systems rely on water as a universal conduit, enabling frequent viral transfer across species. This fluid connectivity broadens the viral host range and may accelerate viral evolution within aquaculture ecosystems.

Given the crucial role of water in viral transmission, effective management of aquaculture water is essential for disease prevention. Many pathogenic shrimp viruses — such as White spot syndrome virus (WSSV), Infectious hypodermal and hematopoietic necrosis virus (IHHNV), and Infectious myonecrosis virus (IMNV) — are known to spread through water^34^. Practices that employ groundwater or disinfected water significantly reduce infection risks, whereas systems relying on untreated seawater experience more frequent outbreaks^27,35^. Importantly, our results reinforce that waterborne transmission extends beyond known pathogens to a wide range of novel viruses, underscoring the need for regular monitoring of aquaculture water using metagenomic approaches.

While our study offers new insights into the aquatic viral diversity and transmission, some limitations remain. The work focused on shrimp monoculture and shrimp-fish polyculture systems within a single geographic region, limiting generalization to other aquaculture models and locations. Future research should encompass diverse culture systems and regional comparisons to better understand ecological and environmental factors influencing viral diversity and transmission.

In conclusion, this study reveals that water not only harbors extensive RNA viral diversity but also shapes the virome composition of aquatic animals. Water acts as both a reservoir and a vector for viral exchange, facilitating cross-species transmission among co-cultured species. These findings highlight the need for integrated water-quality management and virome surveillance to predict, prevent, and mitigate viral outbreaks, ultimately promoting sustainability and resilient aquaculture.

## Methods

### Sample collection

A longitudinal sampling was conducted in shrimp monoculture and shrimp–fish polyculture systems in Maoming, Guangdong Province, China (21.55°N, 111.37°E), from 6 June to 27 September 2020. Ten aquaculture ponds of similar size (∼3300 m^2^; depth ≈ 1.2 m) and management conditions were selected, with five assigned to monoculture and five to polyculture. Sampling in this study encompassed Pacific white shrimp (*Litopenaeus vannamei*), tilapia (*Oreochromis niloticus × O. aureus*), red drum (Sciaenops ocellatus), and aquaculture water.

Sampling covered the entire shrimp culture cycle, including a baseline before larval stocking (Stage 0) and four subsequent stages—early (Stages 1–2), middle (Stage 3), and late (Stage 4)—corresponding to 22, 37, 66, and 107 days after stocking, respectively. Due to high initial salinity, polyculture was initiated after Stage 2, when fish were introduced into ponds containing shrimp from the monoculture group. Shrimp growth cycles were consistent across both systems (Supplementary Fig. 1).

At each stage, 15–20 shrimp per pond were surface sterilized using PBS and dissected (gills, hepatopancreas, and muscle tissues). Ten individuals per fish species were sampled from each polyculture pond at Stage 4. Liver, spleen, kidney, gill, and muscle tissues were dissected. All tissue samples were immediately stored in RNAlater at −80°C. For each pond, 25 liters of water (0.5 m depth) was filtered through a 0.22 μm polyethersulfone membrane, concentrated using tangential flow filtration (50 kDa cutoff) to less than 15 mL, and stored at −80 °C. In total, 525 shrimp, 60 tilapia, 60 red drum, and 42 water samples were collected (Table S1).

### Sample processing and sequencing

Tissue samples were pooled by animal type (shrimp, tilapia, and red drum), culture system (monoculture and polyculture), and culture stage (Stage 0 to Stage 4), yielding 31 shrimp and 12 fish pools. Each pool (15-20 shrimp or 10 fish). Shrimp were processed similar to the procedures used in the previous study^19^. Briefly, tissues were pooled in equal weight ( ∼1 g), homogenized (10% w/v PBS, 60 Hz, 4 °C, 3 min), freeze–thawed three times, and centrifuged (12,000 × g, 10 min). Supernatants were filtered (0.45 μm) and ultracentrifuged (180,000 × g, 3 h). Viral RNA was extracted using the QIAamp Viral RNA Mini kit (QIAGEN, Germany) following the manufacturer’s protocol. Water samples were processed similarly. RNA concentrations were quantified using Qubit 4.0 (Thermo Fisher, USA). Sterile water served as a negative control^36^. Ribosomal RNA (rRNA) was depleted using the NEBNext rRNA Depletion Kit (New England Biolabs, USA), and sequencing libraries were prepared using the NEBNext Ultra RNA Library Prep Kit for Illumina (NEB, USA). These libraries were sequenced on the Illumina NovaSeq platform with 150 bp paired-end reads at Novogene (Beijing, China). One polyculture water sample failed to yield sufficient RNA and was excluded. In total, 84 libraries were sequenced (31 shrimp, 6 tilapia, 6 red drum, and 41 water; Table S1).

### Virus discovery

For each library, adaptor sequences and low-quality bases were trimmed from the raw reads using fastp (v0.21.0)^37^ with the following parameters (SLIDINGWINDOW:4:5, LEADING:5, TRAILING:5, and MINLEN:25). Reads shorter than 100 bp were discarded, and the remaining high-quality reads were de novo assembled into contigs using SPAdes (v3.15.4, “rnaviralspades” module)^38^. Contigs shorter than 600 bp were discarded, and redundancy was reduced using CD-HIT (v4.8.1, 95% nucleotide identity and 80% alignment coverage)^39^.

Viral contigs were via DIAMOND BLASTx (v2.0.11)^40^ against the NCBI non-redundant protein (nr) and the RdRP-scan database^41^ (E < 1e^−5^, very sensitive mode). Candidate viral contigs were further screened with hmmscan (HMMER, v3.3.2)^42^ against two curated RdRP HMM profiles^5,41^ and validated if at least two conserved RdRp motifs were present. Contigs sharing < 90% RdRp protein identity or < 80% nucleotide identity with known viruses were defined as novel species^43^.

Potential host-derived sequences were removed by aligning against shrimp, tilapia, and red drum genomes and the NCBI non-redundant nucleotide (nt) database (blastn v2.13.0). Non-viral contigs meeting ≥ 80% identity over ≥ 50% coverage (or ≥ 90% identity over ≥ 10% coverage) were discarded. Redundant viral sequences were clustered and merged using Geneious Prime (v2023.2.1). No viral sequences were detected in negative controls.

### RNA quantification and quality control

All quality-filtered reads were mapped against the SILVA rRNA database (https://www.arb-silva.de/) using SortMeRNA (v4.3.342)^44^, and non-rRNA reads were retained. These reads were mapped to viral contigs using bwa-mem2 (v2.2.1)^45^ with default settings, and coverage was calculated by CoverM (v.0.6.1)^46^. Specifically, reads mapped at ≥ 90% identity over ≥ 75% of the read length were retained^5^. Viral contigs with horizontal coverage ≥ 30% or length horizontally ≥ 1kb were considered present in a library, and only contigs with horizontal coverage ≥ 99% in at least one library were kept^5^. To remove index-hopping artifacts, any viral read count ≤ 0.1% of the maximum lane abundance was discarded^47^. Viral abundance was normalized as reads per million non-rRNA reads (RPM), with a detection threshold of ≥ 1 RPM.

### Phylogenetic analysis

RdRp sequences from all detected viruses and representative ICTV species were aligned using MAFFT (v7.450, E-INS-i algorithm)^48^, manually refined, and trimmed with trimAl (v1.4.rev15)^49^. Maximum-likelihood phylogenetic trees were inferred using IQ-TREE (v1.6.12, -m MFP)^50^. Branch support was accessed using an approximate likelihood ratio test with Shimodaira-Hasegawa-like (SH-aLRT). Trees were visualized in R (v4.3.1) using the ggtree package (v3.8.0).

### PCR confirmation of high-abundance viruses

High-abundance viruses (> 1000 RPM in shrimp) were validated by nested RT-PCR using specific primers designed from viral contigs. Negative controls used RNase-free water. PCR products were visualized on 1.5% agarose gels and confirmed by Sanger sequencing.

### Statistical methods

Statistical analyses were conducted in R (v4.3.1). Viral alpha diversity (abundance, richness, Shannon index) was calculated using vegan. Between-group differences were tested by the Wilcoxon rank-sum test or Kruskal–Wallis test (ggpubr v0.4.0). Beta diversity was estimated from Bray–Curtis distance matrices and visualized via NMDS (vegan). PERMANOVA (“adonis2”) assessed the effects of sample type, culture system, and culture stage (9999 permutations). Correlations between water and host viromes were analyzed using Spearman’s rank correlation and visualized with ggplot2. Virus-sharing patterns among shrimp, tilapia, and red drum were visualized using E Venn (http://www.ehbio.com/test/venn/#/)^51^ and ComplexHeatmap (v1.0.12).

## Data availability

All raw sequence reads accessible in the SRA databases under BioProject accession XXX. The assembled viral genome sequences have been deposited in the NCBI GenBank database under accession (XXX to XXX).

## Acknowledgements

This study was supported by the National Key Research and Development Program of China (2024YFD2401202), the earmarked fund of China Agriculture Research System for CARS-48 and Southern Marine Science, and Engineering Guangdong Laboratory (Zhuhai) (SML2024SP003).

## Author contributions

J.G.H. and S.P.W. conceived and designed the study; D.D.Z., S.P.W., S.S.L., Z.L.,Y.Q.W., H.L.S., D.R.X., G.Y.G., Z.X.D., S.Z.Z., J.H.P., and Q.L. collected the samples; D.D.Z., Z.L., W.Q.W., H.L.S., D.R.X., G.Y.G., J.R.C., M.H.X., M.C., P.Y.H., C.C.W., and X.Y.H. processed the samples and extracted nucleic acid; D.D.Z. prepared sequencing libraries; D.D.Z., C.G.X., M.S., and J.M.Z. performed the data analysis and visualization; Y.X.L., C.Z., T.Y.W., C.C.Z., Y.C.X., X.C.T., and S.Q.Z. performed the PCR validation; D.D.Z. and C.G.X. wrote the first draft of the manuscript; J.G.H. and M.S. revised the manuscript. All authors reviewed, edited, and approved the manuscript.

