## Supplementary figures S1 ∼ S6 for "RNA viruses in water shape the viromes of shrimp and fish in aquaculture systems"

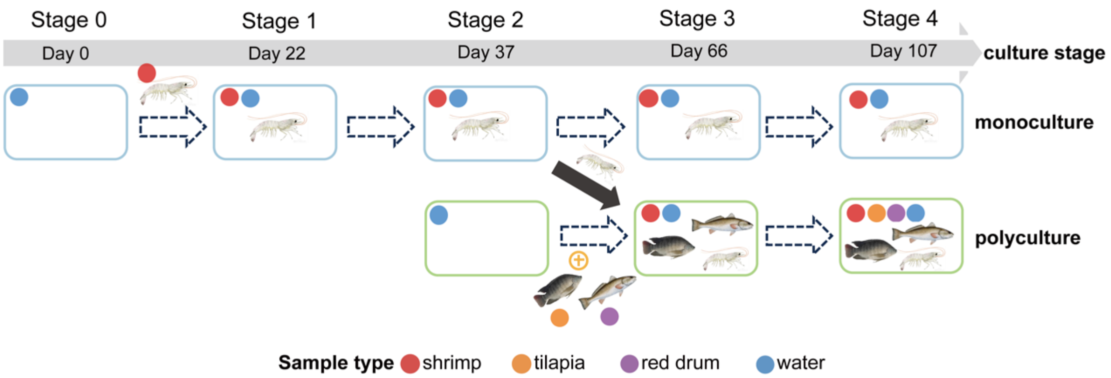


**Supplementary Fig. 1 | Study design and sampling strategy.** This study involved two shrimp culture systems: shrimp monoculture and shrimp-fish polyculture. Sampling was conducted at five key time points: prior to shrimp larval stocking (Stage 0, Day 0) and at 22, 37, 66, and 107 days after the start of the culture cycle (Stages 1 to 4). Stage 1 and Stage 2 represent the early growth period of shrimp, Stage 3 represents the middle stage of shrimp farming, and Stage 4 corresponds to the late stage of the shrimp culture cycle, just prior to harvest. The monoculture system was sampled across all five stages, while the polyculture system was sampled starting from Stage 2. Circles indicate the types of samples collected at each time point: red for shrimp, orange for tilapia, purple for red drum, and blue for water. The arrow indicates the transfer of shrimp from the monoculture system to the polyculture system after Stage 2.

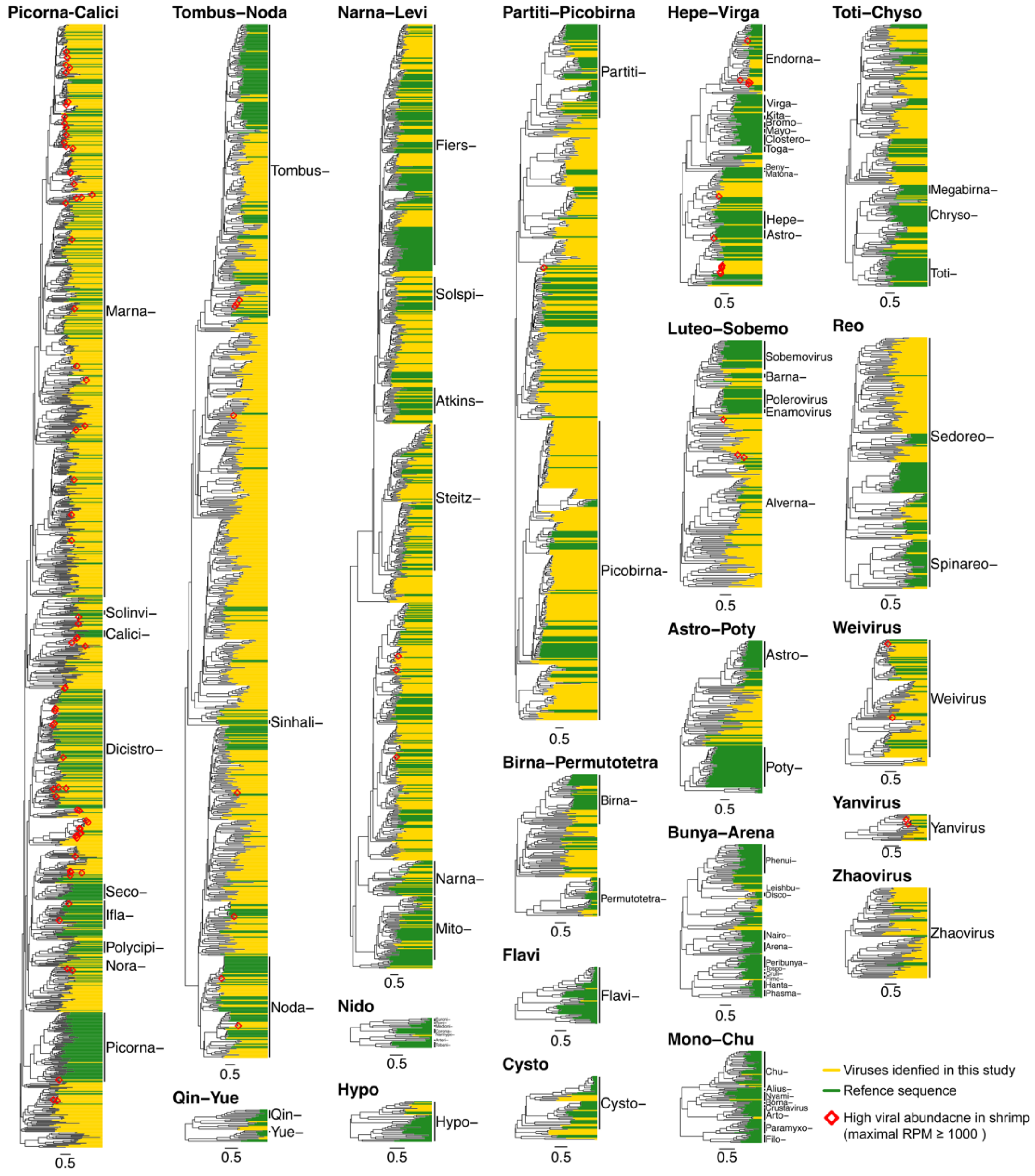


**Supplementary Fig. 2 | Phylogenetic analysis of the 3211 RNA viruses found in shrimp aquaculture systems.** The maximum likelihood phylogenetic trees were estimated at the ‘supergroup’ level based on RNA-dependent RNA polymerase (RdRp) protein. Vertical lines and abbreviated family names (without the suffix '-viridae') were used to denote viral families on the right side of each tree. Reference sequences are depicted in green, while viruses identified here are denoted in yellow. Viruses that exhibited high abundance (reads per million total reads (RPM) ≥1000) in shrimp are denoted by red diamonds. Trees are midpoint rooted for clarity, and branch lengths are indicated by the scale bar. Each scale bar indicates 0.5 amino acid substitution per site.

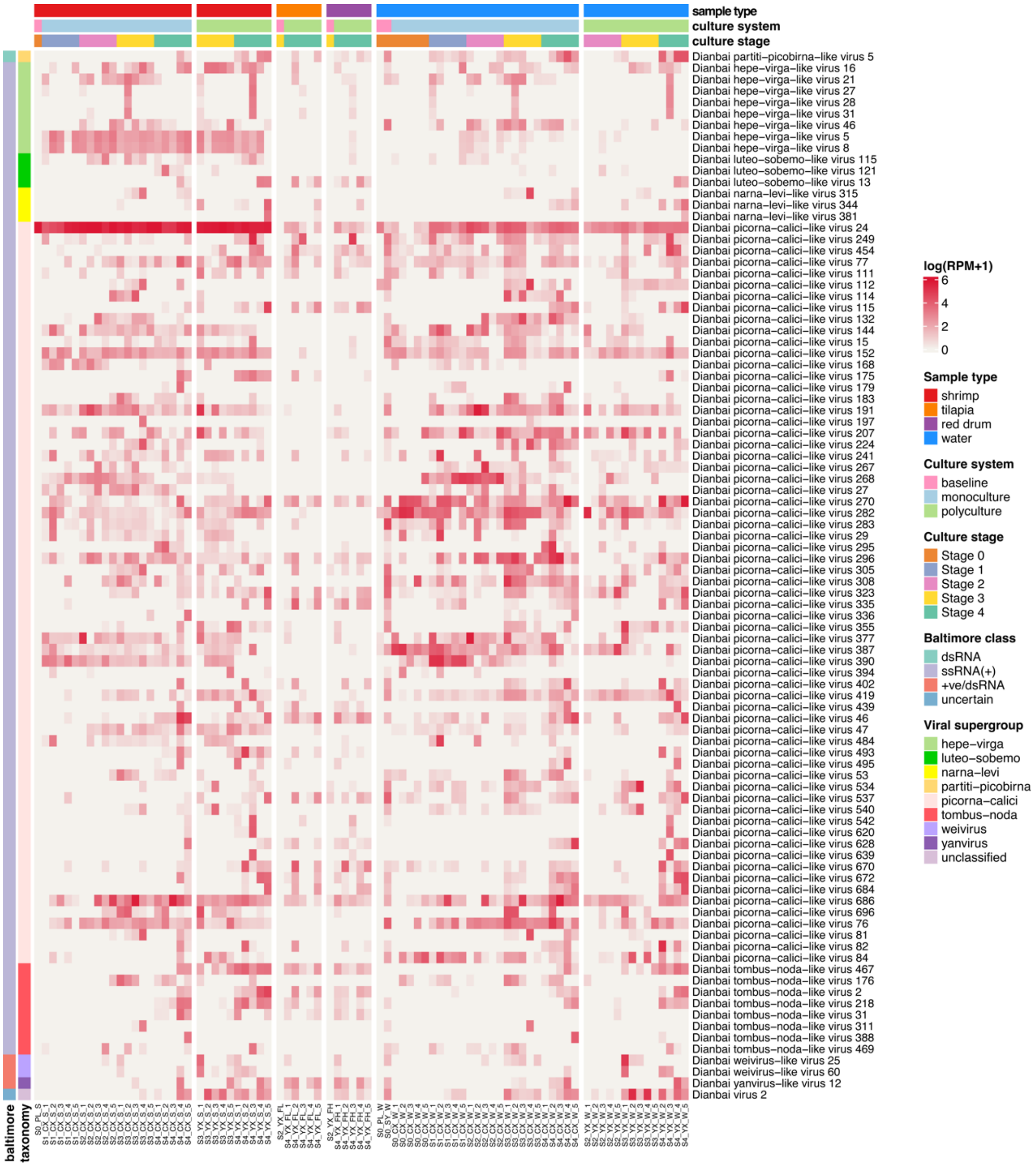


**Supplementary Fig. 3 | The heatmap illustrates the distribution and abundance of high-abundance viruses in each library.** The abundance of viruses in each library was normalized by a logarithm of RPM. Each column represents a library, while each row represents a virus species. Sample types, culture systems, and culture stages are shown as colored strips at the top. The viral supergroups and baltimore class of viruses are shown on the left. Samples (x-axis) are divided based on sample types and culture systems into six different groups. Virus species (y-axis) are ordered based on viral taxonomy, including eight viral supergroups and one unclassified virus.

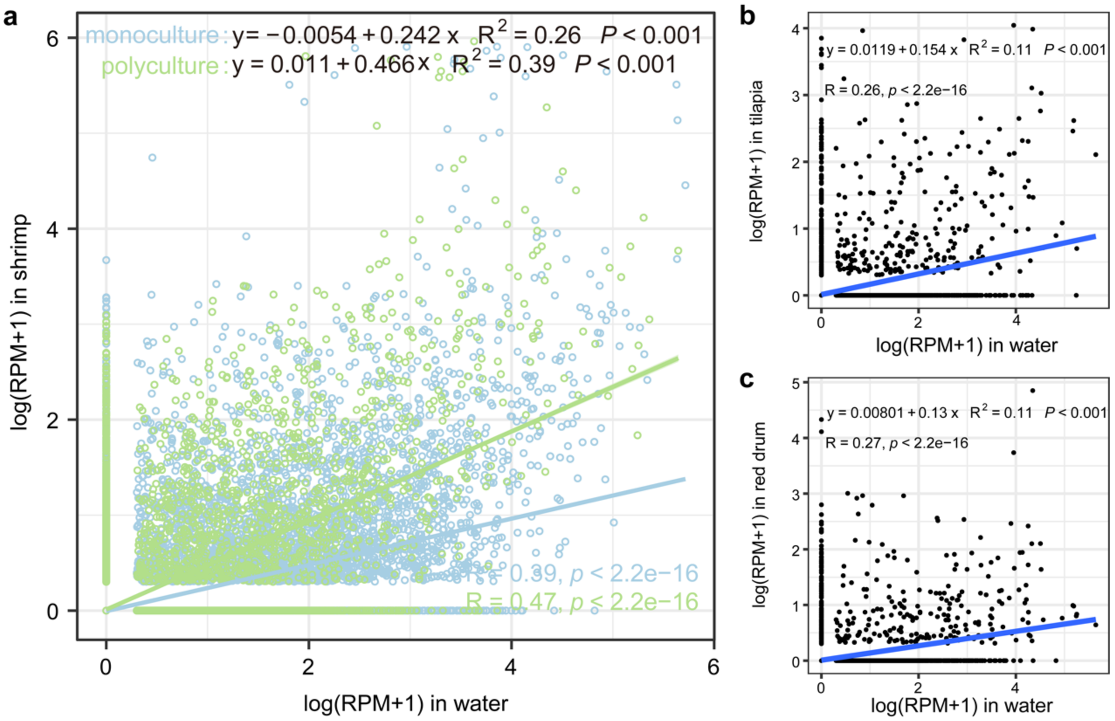


**Supplementary Fig. 4 | Linear regression showing the correlation of the abundance of 3211 viruses between aquatic animals and water.**

**a**, Correlation of viral abundance between shrimp and water. Shaded areas represent 95% confidence intervals.

**b**, Correlation between tilapia and water.

**c**, Correlation between red drum and water.

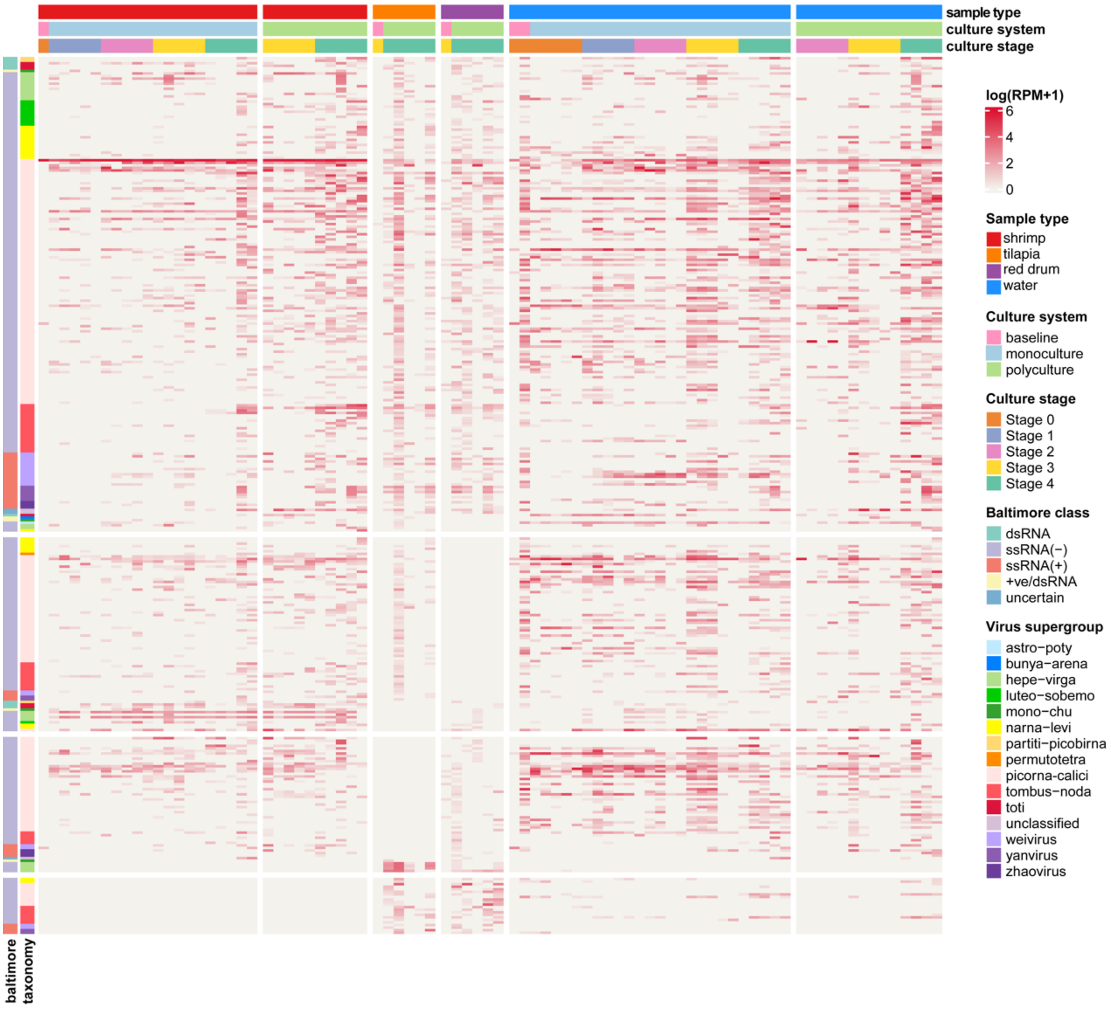


**Supplementary Fig. 5 | The heatmap illustrates the distribution and abundance of virus-sharing species in each library.** The abundance of viruses in each library was normalized by a logarithm of RPM. Each column represents a library, while each row represents a virus species. Sample types, culture systems, and culture stages are shown as colored strips at the top. The viral supergroups and baltimore class of viruses are shown on the left. Samples (x-axis) are divided based on sample types and culture systems into six different groups, and virus species (y-axis) are ordered based on viral taxonomy.

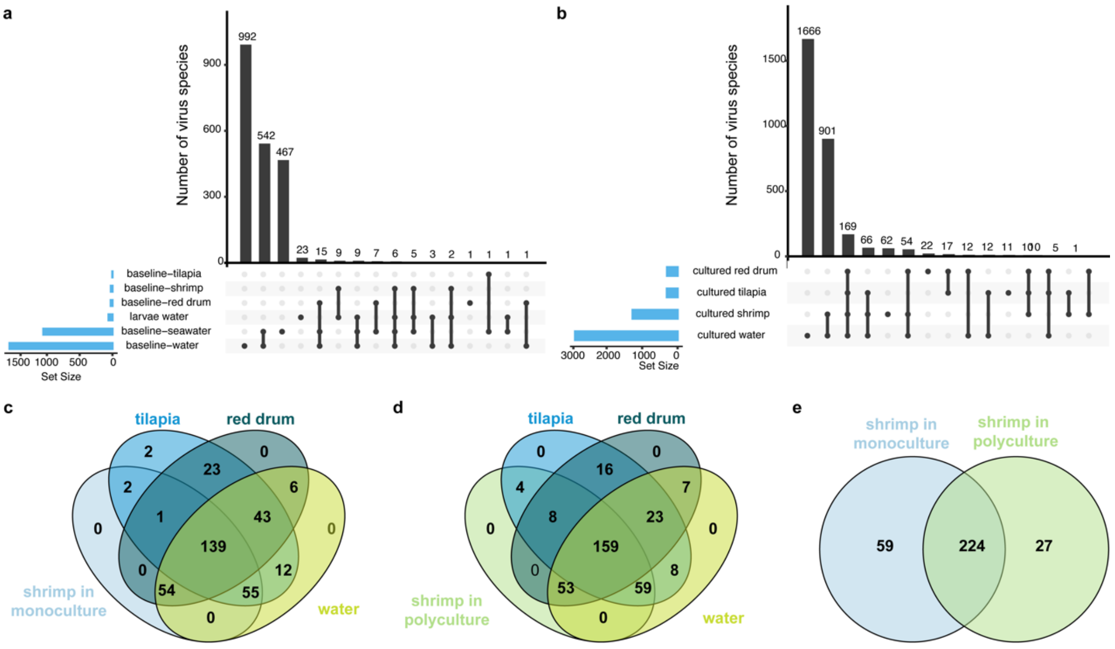


**Supplementary Fig. 6 | a**, Upset diagram displaying the overlap of virus species among different baseline samples.

**b**, Overlap of virus species among different samples during culture period.

**c**, Venn diagram showing the overlap of virus-sharing species among shrimp from the monoculture system, tilapia, red drum, and water.

**d**, Overlap of virus-sharing species among shrimp from the polyculture system, tilapia, red drum, and water.

**e**, Overlap of virus-sharing species in shrimp between two culture systems.
